# Cooperative self-assembly of nanoparticle-encapsulating hybrid protein cages

**DOI:** 10.64898/2026.01.21.700743

**Authors:** Suna Jo, Won Min Park

## Abstract

Protein cages are versatile platforms capable of encapsulating a wide range of nanoparticle cargo within biocompatible protein shells while providing tunable functionalities. Here, we investigated a self-assembly system that forms vesicle-like protein cages while simultaneously encapsulating nanoparticles at high density, yielding pomegranate-like protein– nanoparticle hybrid materials. Amphiphilic recombinant fusion protein building blocks based on elastin-like polypeptides, leucin zippers, and fluorescent proteins were employed to assemble vesicle-like protein cages via temperature-triggered liquid-liquid phase separation in the presence of fluorescent polystyrene nanoparticles. Analysis of nanoparticle encapsulation density and protein cage size indicates cooperative interactions between protein building blocks and nanoparticles that mediate the formation of protein-nanoparticle coacervate intermediates, which subsequently convert into core-shell hybrid protein cages, as further supported by kinetics studies. We demonstrate the self-assembly hybrid protein cages incorporating a fluorescent calcium sensor protein and titanium oxide nanoparticles, which exhibit a drastic enhancement in their calcium-sensing capability as a result of nanoparticle encapsulation. This platform offers a broadly applicable strategy that integrates protein biofunctionality with diverse nanoparticle properties for development of advanced hybrid materials.

## 1. Introduction

Protein-based materials have been extensively investigated as hybrid materials in combination with non-protein foreign components due to their ability to form diverse structures and control materials’ properties.^[1–4]^ These hybrid materials improve the physical and chemical properties of the protein materials, making them more promising. Specifically, protein-based hybrid materials have advanced biotechnology, including biocatalysts^[5]^, biosensors^[6]^, and targeted drug delivery systems.^[7,8]^ Proteins can be engineered into various structures, which serve as templates to improve material solubility^[9]^, stability^[10]^, and biocompatibility.^[11–13]^

Among the structures found in protein-based materials, protein cages (PCs) are of particular interest. They feature a spherical morphology with an inner cavity capable of capturing molecules while isolating them from the external environment.^[14,15]^ This feature facilitates the incorporation of foreign materials within a confined space.^[16– 19]^ Recent studies have demonstrated that nanoparticles (NPs) can be encapsulated within the inner cavity, while the exterior surface retains favorable biocompatibility and targeting capabilities.^[10,20–22]^ Consequently, PCs have attracted growing attention for drug delivery applications, as their distinctive morphology and size support efficient drug encapsulation and cellular delivery.^[18,23–28]^

A PC typically consists of three interfaces: the exterior surface, the inter-subunit region, and the inner cavity. ^[29–32]^ The properties of PCs can be modulated by incorporating foreign elements for high functional versatility. For instance, surface modification of the exterior can introduce cell-targeting ligands for a therapeutic application. The inner cavity can accommodate functional NPs, such as quantum dot^[33]^, gold^[30]^, and iron oxide.^[34]^ Ferritin, for example, has been utilized as a template for synthesizing or encapsulating inorganic NPs for cancer cell imaging.^[30,35]^ Similarly, virus-like particles have been chemically modified to load and deliver therapeutic cargo.^[36,37]^

These PCs often exhibit limited NP loading capacity. To date, high-density NP encapsulation has been achieved in amphiphilic block-copolymer micelles,^[38–40]^ and also in vesicle-like PCs.^[41]^ In the latter example, amphiphilic protein building blocks were engineered using elastin-like polypeptide (ELPs), a class of biomimetic polypeptides derived from tropoelastin.^[42]^ ELPs exhibit lower critical solution temperature phase behavior, which mediates temperature-controlled liquid-liquid phase separation (LLPS) into protein coacervates,^[43–45]^ and function as the hydrophobic segment in amphiphilic protein buillding blocks.^[41]^ ELPs were recombined with globular (mCherry) or rod-shaped heterodimeric leucine zipper proteins (Z_R_ and Z_E_) to generate two recombinant fusion proteins, mCherry-Z_E_ and Z_R_-ELP (**Figure 1**). Upon mixing, these fusion proteins form “globule-coil-rod” (mCherry-Z_E_/Z_R_-ELP) or “rod-coil” (Z_R_-ELP dimer) amphiphilic complexes, which further assemble into vesicle-like hollow PCs upon temperature elevation.^[41]^

**Figure 1.**
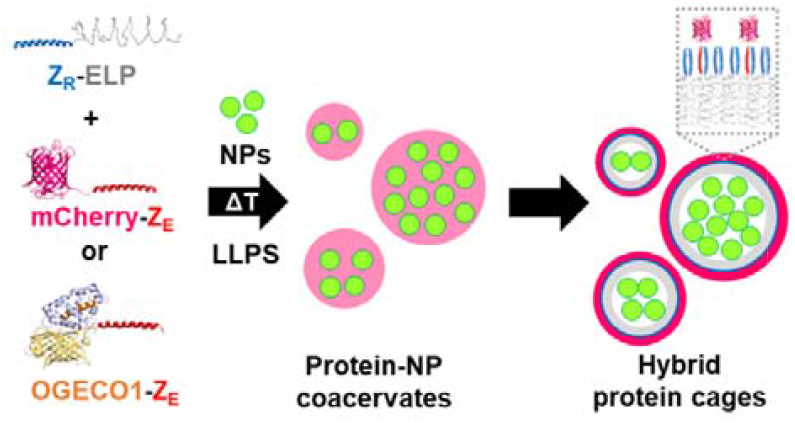
Cooperative self-assembly of hybrid PCs encapsulating NPs. Amphiphilic protein complexes from the recombinant fusion proteins Z_R_-ELP and mCherry-Z_E_ (or OGECO1-Z_E_) have cooperative interactions with NPs, forming a protein-NP hybrid coacervate that further evolves into hybrid PCs with high-density packing of NPs.

The resulting PCs consist of a monolayer protein membrane with hydrophobic inner surfaces.^[41]^ This unique architecture facilitates preferential encapsulation of NPs within the PC, resulting in pomegranate-like structures with a high-density packing of NPs.^[41]^ Despite this remarkable ability to encapsulate NP cargo, the mechanism underlying preferential and high-loading NP encapsulation during PC assembly remains unclear. We hypothesize that an intermediate stage of protein-NP coacervate formation is invovled (**Figure 1**), and to test this, we investigated the copperativity between protein-protein and protein-NP interations that drives to the assembly of NP-encasulated PCs. Using this system, we further demonstrated tunable optical properties of hybrid PCs for fluorescent biosensor applications. Specifically, the red fluorescent calcium sensor protein OGECO1^[46]^ was engineered into a recombinant fusion protein (OGECO1-Z_E_; **Figure 1** and **Figure S1** in Supporting Information) to assemble hybrid PCs.

## 2. Results and Discussion

### 2.1 Vesicle-Like PC Assembly and NP Encapsulation

We prepared the recombinant fusion proteins Z_R_-ELP and mCherry-Z_E_ (**Figure 1**) in phosphate buffered saline (PBS) and mixed them with polystyrene (PS) NPs (diameter ∼ 500 nm). The self-assembly of PCs was induced by increasing the temperature from 4°C to 25°C. This temperature elevation resulted in a rapid increase in turbidity, confirming the formation of vesicle-like PCs. Fluorescent microscope images revealed pomegranate-like morphologies where PS NPs (green) were encapsulated by the self-assembled protein membrane (red) (**Figure 2A**). The protein membrane contained mCherry with red fluorescence, while the PS NPs exhibited green fluorescence. The internal space of the vesicle-like PCs was densely packed with PS NPs, which distinguished them from the morphologies observed in NP-encapsulating PCs reported in other studies.^[30,47,48]^ The number of encapsulated PS NPs (*n*_*NP*_) observed in the microscope images ranged from 1 to over 14 (**Figure 2B**).

**Figure 2.**
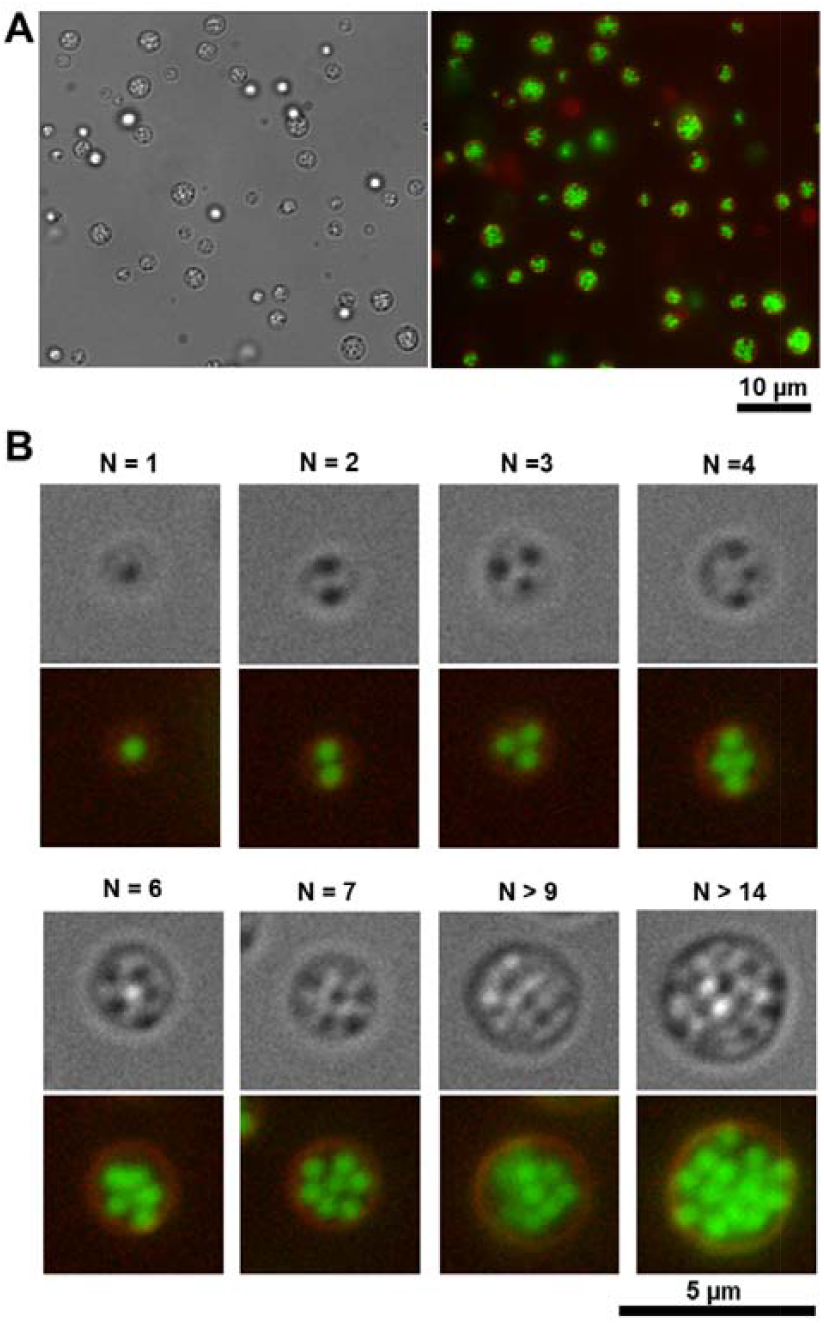
The morphology of PCs encapsulating PS NPs. (A) Fluorescent microscope images. The red fluorescence is attributed to mCherry-Z_E_, while the green fluorescence is from the PS NPs. (B) Individual PCs and the number of PS NPs encapsulated withi**n** each PC.

Next, we estimated the relationship between the size of PCs and encapsulated PS NPs and observed that the diameters of PCs were proportional to the number of encapsulated PS NPs (**Figure 2A**). The average diameter of PCs containing NPs (*d*_*PC*_ ∼ 2.3 ± 0.6 μm; **Figure S2A**) was larger than that of the hollow PCs without NPs (∼1.5 μm),^[41]^ indicating the presence of NPs during PC formation affected the size of resulting PCs. The image analysis demonstrated that the number of PS NPs counted from cross-sectional images was approximately proportional to the cross-section area of the PCs (*n*_*NP*_ ∼ *d*_*PC*2.3_). Consistently, the volume-averaged number of PS NPs encapsulated within each PC (*n*_*NP*_) scales to the volume of PCs (*n*_*NP*_ ∼ *d*_*PC*3_, **Figures 3A. and 3B**). The dependency of PC size on the number of encapsulated NPs supports the existence of cooperative interactions that facilitate the incorporation of NPs during the coacervates formation, ultimately leading to a core-shell morphology. In the absence of NPs, it has been demonstrated that the intermediate coacervate phase transitions into hollow PCs, further supporting this mechanism.^[49,50]^ Accordingly, the estimated volume fraction of PS NPs per PC (V_NP_/V_PC_; **Figure 3B**) showed a narrow distribution, indicating that NP packing density is highly controlled. The number of NPs encapsulated within PCs varies widely, from just a few to over 40 (**Figure S2B**), while the volume fractions of NPs per PC remain consistent.

**Figure 3.**
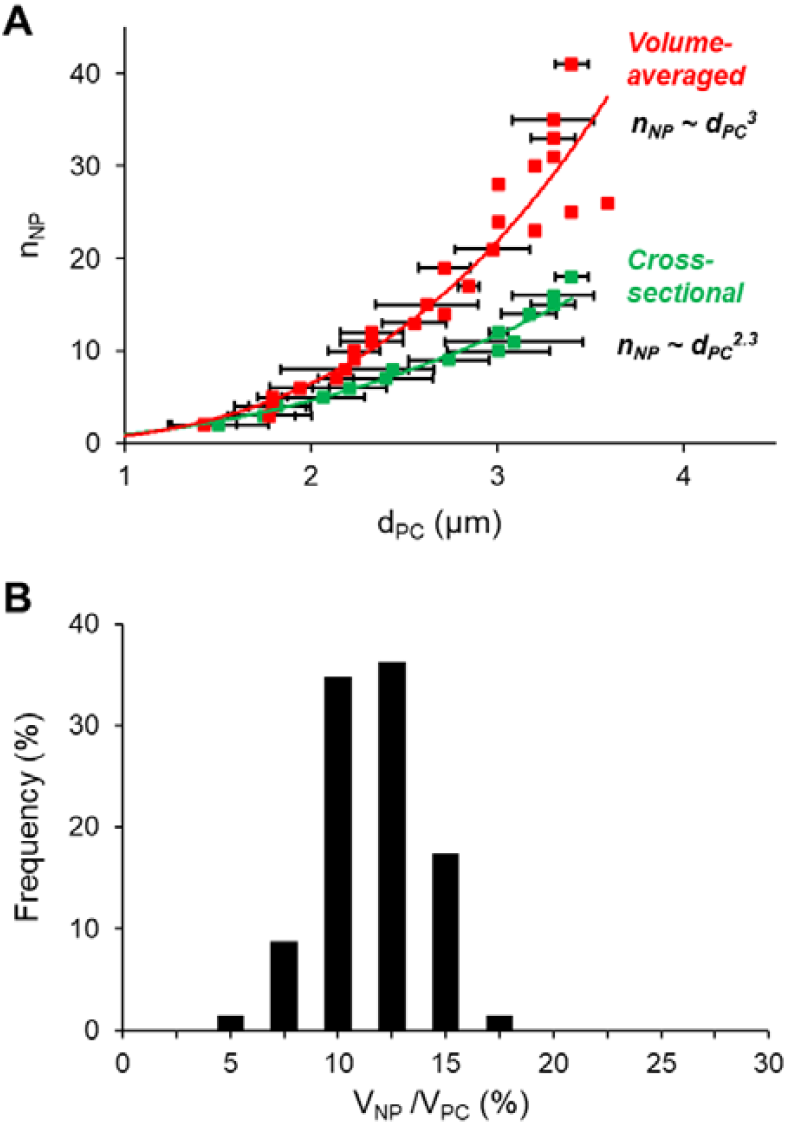
Relationship between PC size and PS NPs. (A) Correlation between the diameter of PCs (*d*_*PC*_) and the number of NPs (n_NP_) counted from cross-sectional areas and estimated from PC volume. (B) Distribution of volume fraction of PS NPs (V_NP_/V_PC_) as a percentage of total frequency.

### 2.2 Kinetics of NP Encapsulation

The turbidity profiles further support the cooperative interactions between the protein building blocks and PS NPs during PC assembly. In the presence of NPs (ranging from 3.64 to 18.2 particles/μL), turbidity changes during the initial inverse phase transition were measured as a function of time. Turbidity is associated with the presence of coacervates.^[41]^ In the initial stage for coacervate formation, turbidity increases rapidly, followed by a gradual decrease during the maturation stage that involves coalescence of protein coacervates. In the presence of NPs, a slower rise in turbidity profile after 4 minutes was observed at higher concentrations of NPs (**Figure. 4A**). To be consistent, a steeper decline in the slope of turbidity change was observed during this period at higher NP concentrations (**Figure. 4B**). The results suggest that coalescence of protein coacervates and PS NPs occur after the initial stage for coacervate formation. According to the mechanism of hollow Z_R_-ELP/mCherry-Z_E_ vesicles,^[50]^ the process involves coalescence of protein coacervates, followed by a transition to hollow vesicles. Similarly, the NP-bound protein coacervates are likely to undergo a transition to vesicles, during which the PS NPs are placed in the core, and the amphiphilic protein complexes organize into a membrane structure that encloses the NPs. This step may appear as an increase in the slope of the turbidity profile (indicated by the arrows in **Figure 4B**), which is delayed in the presence of NPs during PC assembly. This supports that NPs participate in the coacervate formation, leading to a slower transition into the hybrid PCs due to increased interactions with more NPs.

**Figure 4.**
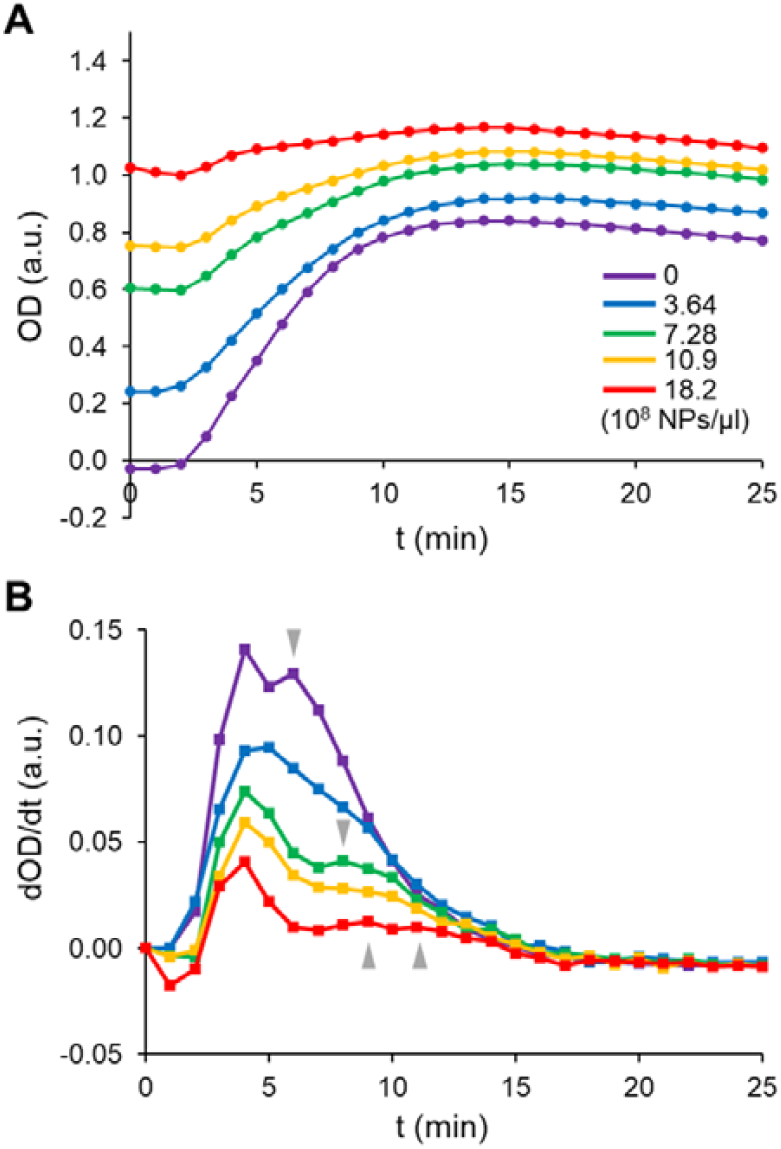
Turbidity changes as a function of time (t) during PC assembly. (A) Optical density (OD) profiles at 400 nm. (B) Changes in the slope of the turbidity profiles (dOD/dt). Samples were initially pre**p**ared at 4°C with varying NP concentrations (purple: no NPs; blue: 3.64 × 10^8^ particles/µl; green: 7.28 × 10^8^ particles/µl; yellow: 10.9 × 10^8^ particles/µl; red: 18.2 × 10^8^ particles/µl) and OD was measured upo**n** heat-ing to 25°C.

### 2.3 Encapsulation of Titanium Dioxide (TiO_2_) NPs

To evaluate whether different types of NPs can be encapsulated, we used this platform for the encapsulation of TiO_2_ NPs. TiO_2_ NPs are well known as a sunscreen agent^[51]^ and a photocatalyst^[52]^ due to their effective absorption and radical generation under UV irradiation. However, TiO_2_ can release reactive oxygen species (ROS), which may promote skin photodamage throphotocatalytic degradation.^[53]^ Surface inhibition of TiO_2_ particles, if encapsulation is successful, could reduce photocatalytic activity. To test this hypothesis, we measured the photocatalytic activity of TiO_2_ encapsulated by PCs under UV radiation.

Hybrid PC assembly was carried out in the presence of TiO_2_ NPs, as confirmed by fluorescence microscopy images showing the formation of fluorescent particles with a core–shell morphology (Figure S3). A photodegradation experiment with TiO_2_ was conducted under various sample conditions, using Rhodamine B (RhB) as a model dye to assess the reaction. RhB is a dye that undergoes discoloration upon reaction with radicals generated by TiO_2_ under UV irradiation. Control samples with TiO_2_ without UV exposure, as well as samples exposed to UV light in the absence of TiO_2_, did not show any photodegradation of RhB. Under UV irradiation, TiO_2_ without proteins exhibits the fastest photocatalytic degradation, whereas samples containing TiO_2_-encapsulated PCs showed no significant photodegradation but a slight increase in turbidity (**Figures 5A** and **S4**). These findings suggest that TiO_2_ NPs were encapsulated by the protein layer, which suppresses radical release even under UV irradiation. We further investigated the UV-screening property of the TiO_2_-encapsulated PCs by measuring the transmittance in the UV range from 200 nm to 300 nm (**Figure 5B**). The hybrid sample exhibited the lowest UV-transmittance compared to the samples containing TiO_2_ or PCs only, indicating superior UV-blocking capabilities.

**Figure 5.**
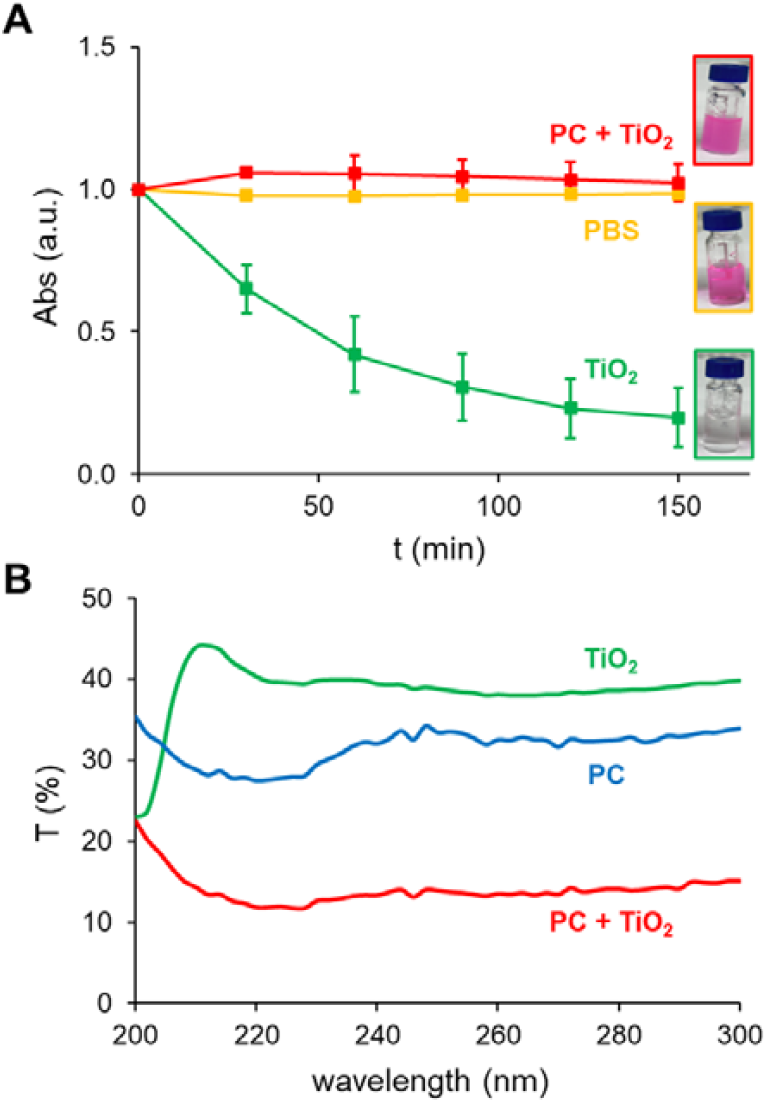
Photocatalytic activity of TiO_2_ NP-encapsulating hybrid PCs. (A) Normalized absorbance (Abs) changes at 554 nm. RhB samples included 2X PBS only (yellow), TiO_2_ under UV irradiation (green), and TiO_2_-encapsulated PCs under UV irradiation (red). (B) Transmittance (T) spectra of TiO_2_ NPs (green), TiO_2_-encapsulated PCs (red), and PCs only (blue). Error bars: standard deviation (n = 3).

### 2.4 Calcium Sensing Using TiO_2_-Encapsulated Hybrid PCs

In addition to strong UV absorption, TiO_2_ is known for its strong light-scattering properties due to its high refractive index. This property allows TiO_2_ to effectively reflect and scatter light,^[54,55]^ which can enhance the fluorescence of molecular agents when combined with TiO_2_.^[56]^ For example, the fluorescence enhancement effect of TiO_2_ in cancer detection surgeries has been demonstrated using polyethylene glycol-modified TiO_2_ NPs in combination with photodynamic diagnosis agents for bladder cancer cells.^[56]^ To explore this property of TiO for biosensor applications, we integrated the red fluorescent calcium sensor protein OGECO1^[46]^ into TiO -encapsulating hybrid PCs.

We designed the recombinant fluorescent fusion protein OGECO1-Z_E_ that can assemble with Z_R_-ELP to form hybrid PCs encapsulating TiO_2_ NPs, and confirmed absorbance and fluorescence change of OGECO1-Z_E_ upon binding to Ca^2+^ (**Figures S1** and **S5**). Hybrid PCs of Z_R_-ELP and OGECO1-Z_E_ (with an OGECO1-Z_E_:Z_R_-ELP ratio of 0.05) were prepared at varying free Ca^2+^ concentrations, and fluorescence amplification relative to the no Ca^2+^ condition (ΔF/F_0_) was measured. Upon encapsulation of TiO_2_ NPs, a substantial enhancement in the fluorescence amplification (> 2500 %) was observed (**Figure 6A**). Specifically, at a free Ca^2+^ concentration of 0.78 nM, PCs without TiO_2_ exhibited a ΔF/F_0_ of 262.9 (± 26.7), while the inclusion of TiO_2_ increased this value to 850.7 (± 144.4), corresponding to more than a threefold enhancement (**Figure 6B**). This pronounced increase in fluorescence intensity and sensitivity, facilitated by TiO_2_ encapsulation, highlights the potential of this system as a platform for future biosensor applications.

**Figure 6.**
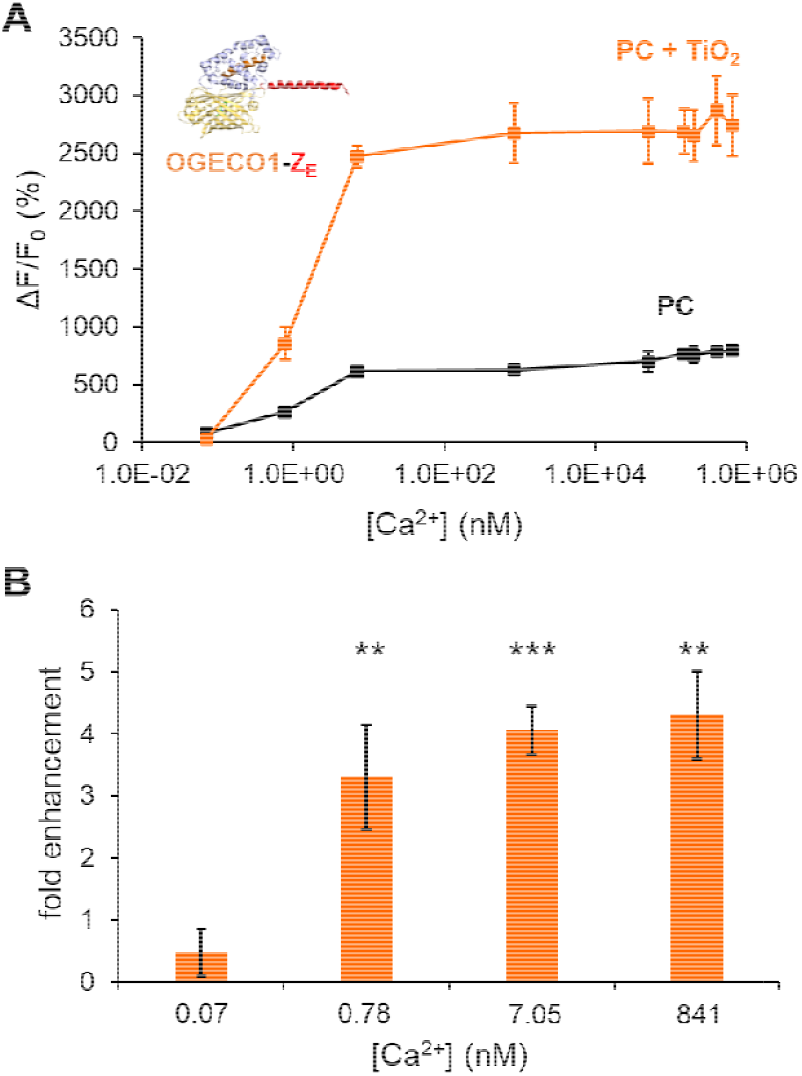
Enhanced calcium sensing using PCs (OGECO1-Z_E_/Z_R_-ELP) encapsulating TiO_2_ NPs. (A) Relative fluorescent amplification (ΔF/F_0_) as a function of free Ca^2+^ concentrations. (B) Fold enhancement in ΔF/F_0_ by TiO_2_ NP encapsulation compared to PCs only, across different concentrations of free Ca^2+^: 0.07 nM, 0.78 nM, 7.05 nM, 841 nM. **p ≤ 0.01 and ***p ≤ 0.001. Error bars: standard deviation (n = 3).

## 3. Conclusion

We investigated the cooperative self-assembly of PCs capable of simultaneous encapsulation of NPs with highdensity packing. Our analysis using fluorescent PS NPs supports the cooperative interactions between protein building blocks and NPs during the formation of protein-NP coacervate intermediates, which subsequently converted into core-shell hybrid PCs. These findings were further supported by a kinetics study. This versatile platform was extended to TiO_2_ NPs, where suppressed photocatalytic activity further confirmed the formation of a protein layer encapsulating NPs. Furthermore, TiO_2_-encapsulating PCs exhibited significant fluorescence enhancement when integrated with fluorescent protein calcium sensor probes. This modular and versatile platform can accommodate a range of NPs and fluorescent protein probes, offering potential applications in nanomedicine, biosensors, and bioimaging.

## 4. Materials and Methods

### 4.1 Materials

The E. coli strain T7 Express BL 21 was purchased from New England Biolabs. The plasmids, pQE60_Z_E_/Z_R_-ELP and pQE60_mCherry−Z_E_, were also kindly provided by Professor Julie Champion at the Georgia Institute of Technology. Nickel nitrilotriacetic acid (HisPur™ Ni-NTA) resin was purchased from Thermo Scientific. Fluorescent polystyrene NPs (Fluoresbrite® YG Carboxylate Microspheres 0.50 μm) were purchased from Polysciences. TiO_2_ NPs, the mixture of rutile and anatase (d < 100nm), were purchased from Sigma Aldrich. RhB was purchased from ACROS Organics. PBS (1X at pH 7.4) containing 137 mM NaCl, 2.7 mM KCl, 10 mM NaH_2_PO_4_, 1.8 mM KH_2_PO_4_ (1.8 mM), as well as tris buffered saline (TBS at pH 7.4) containing 50 mM tris base and 150 mM NaCl were prepared in the lab.

#### 4.2 Cloning, Protein Expression, and Purification

The DNA sequences encoding OGEO1^[46]^ and the Z_E_ domain^[57]^ were combined to generate a DNA construct encoding OGECO1-Z_E_. The gene fragment was synthesized and inserted into the *NcoI* and *BglII* restriction sites of the pQE60 vector (3.4kbp), followed by sequence verification by GenScript. The resulting plasmid pQE60_OGECO1-Z_E_ was inserted into T7 Express cells. The fusion proteins Z_R_-ELP, mCherry-Z_E_, and OGECO1-Z_E_ were expressed overnight and purified via metal affinity chromatography using hexa-histidine tag (His_6_), following previously described protocols.^[41,58]^ Briefly, Z_R_-ELP was co-expressed with His_6_-Z_E_ and incubated with Ni-NTA after cell harvesting and lysis using a buffer containing 8 M urea, 10 mM Tris-Cl, and 100 mM Na_2_HPO_4_ at pH 8.0. 6 M guanidine hydrochloride was used to elute Z_R_-ELP after washing, then dialyzed in deionized water and stored at 4°C. The cells expressing mCherry-Z_E_ and OGECO1-Z_E_ were lysed with a buffer containing 50 mM Na_2_HPO_4_, 300 mM NaCl, and 10 mM imidazole, and the proteins were purified under native conditions using Ni-NTA by elution with a buffer containing 250 mM imidazole after washing. The eluted fractions were dialyzed in the 1X TBS.

#### 4.3 Fluorescence Microscopy

The fusion protein mixture containing 30 μM Z_R_-ELP, 1.5 μM mCherry-Z_E_ was prepared, to which PS NPs were added at a concentration of 18.2 × 10^8^ NPs/µl and at 4°C, followed by incubation at room temperature for 30 minutes. For the hybrid PCs with TiO_2_ NPs, a TiO_2_ suspension was added to the same protein mixture above at a final concentration of 0.25% (w/v). Images of the hybrid PCs were obtained with an inverted fluorescent microscopy (Axio Observer.Z1 and Olympus IX73). A 5 µl droplet of the sample was placed on a slide glass (24 x 60 mm), covered with a square cover glass (18 x 18 mm). All images are captured with 40x or 100x objectives.

#### 4.4 Turbidity and UV-Transmittance Measurement

The fusion protein solutions were prepared by mixing Z_R_-ELP (30 μM) and mCherry-Z_E_ (1.5 μM) at 4°C, followed by adding fluorescent PS NP solutions at different total NP concentrations. After transferring to a cuvette, the OD at 400 nm was monitored for 30 minutes at 25°C using a spectrophotometer (Thermo Fisher BIOMATE 160), with data collected at 1-minute intervals. For transmittance, samples containing both proteins and 0.25% (w/v) TiO_2_, proteins without TiO_2_, or TiO_2_ without proteins were prepared at 4°C, followed by incubation at 25°C and spectral acquisition from 200 nm to 300 nm using a 1 mm quartz cuvette.

#### 4.5 Photocatalytic Activity Measurement

All samples contained 20 μM RhB in 2X PBS. Samples containing proteins (30 μM Z_R_-ELP and 1.5 μM mCherry-Z_E_), TiO_2_ NPs, or both were prepared in a 1 mL glass vial and incubated at room temperature for 30 minutes. Following incubation, samples were exposed to UV irradiation (BTLab Systems, 302 nm, 50–60 Hz), and the absorbance at 554 nm was measured at hourly intervals to monitor photocatalytic activity.

#### 4.6 Fluorescent Sensing of Calcium Ions

Fluorescence measurements were performed at 25 °C using a 96-well microplate reader (Biotek) for samples containing PCs (30 μM Z_R_-ELP and 1.5 μM OGECO1-Z_E_) with or without 0.125% (w/v) TiO_2_. Fluorescence was recorded with an excitation wavelength of 520 nm and emission at 570nm. To achieve a Ca^2+^-free state, OGECO1-Z_E_ was treated with 100 µM of ethylenediaminetetraacetic acid (EDTA). CaCl_2_ solution was then added to achieve varying final concentrations of free Ca^2+^. Free Ca^2+^ concentrations in the presence of EDTA were calculated using the Maxchelator tool.^[59]^

## Acknowledgements

This research was funded by grants from the National Science Foundation (2239927), in part by the Johnson Cancer Research Center at Kansas State University, and by the Grants4Ag program from Bayer Crop Science. We acknowledge Prof. Julie Champion (Georgia Institute of Technology) for plasmid DNA and Tej Shrestha at the Nanotechnology Core in the College of Veterinary Medicine (Kansas State University) for the assistance in microscope imaging.

## Author Contributions

W.M.P. conceptualized the project. S.J. and W.M.P. designed proteins, planned and performed experiments, interpreted the data, and wrote the manuscript. W.M.P. oversaw all aspects of the project.

## Note

This manuscript is partially derived from a chapter of one author’s doctoral thesis completed at Kansas State University (2024).

