## Supporting Information for "Cooperative self-assembly of nanoparticle-encapsulating hybrid protein cages"

**Table S1.** Amino acid sequences of the recombinant protein building blocks, $Z_R$-ELP, mCherry-$Z_E$, and OGECO1-$Z_E$. The protein domains and motifs are colored: $Z_R$ (blue), $Z_E$ (red), ELP (gray), mCherry (pink), and OGCEO1 (orange).

| Protein | Sequence |
|---|---|
| $Z_R$-ELP | MKGSLEIRAAALRRRNTALRTRVAELRQRVQRLRNEVSQYETRYGPL$(G_4S)_2$G$[(VPGVG)_2VPGFG(VPGVG)_2]_5$VPGC |
| mCherry-$Z_E$ | MGGSRSMVSKGEEDNMAIIKEFMRFKVHMEGSVNGHEFEIEGEGEGRPYEGTQTAKLKVTKGGPLPFAWDILSPQFMYGSKAYVKHPADIPDYLKLSFPEGFKWERVMNFEDGGVVTVTQDSSLQDGEFIYKVKLRGTNFPSDGPVMQKKTMGWEASSERMYPEDGALKGEIKQRLKLKDGGHYDAEVKTTYKAKKPVQLPGAYNVNIKLDITSHNEDYTIVEQYERAEGRHSTGGMDELYKSKLRGSGSLEIEAAALEQENTALETEVAELEQEVQRLENIVSQYRTRYGPLRSHHHHHH |
| OGECO1-$Z_E$ | MVDSSRRKWIKAGHAVRAIGRLSSPVVSERMYPEDGVLKSEIKKGLRLKDGGHYAAEVKTTYKAKKPVQLPGAYIVDIKLDIVSHNEDYTIVEQCERAEGRHPTGGRDELYKGGTGGSLVSKGEEDNMAIIKEFMRFKVHMEGSVNGHEFEIEGEGEGRPYEAFQTAKLKVTKGGPLPFAWDILSPQFTYGSKAYIKHPADIPDYFKLSFPEGFRWERVMNFEDGGIIHVNQDSSLQDGVFIYKVKLRGTNFPPDGPVMQKKTMGWEATRDQLTEEQIAEFKEAFSLFDKDGDGTITTKELGTVLRSLGQNPTEAELQDMINEVDADGDGTFDFPEFLTMMARRMNDTDSEVEIREAFRVFDNDGNGYIGAAELRHVMTDLGEKLTDEEVDEMIRVADIDGDGQVNYEEFVQMMTAKGGSGGTGGSGGTGGSLEIEAAALEQENTALETEVAELEQEVQRLENIVSQYRTRYGPLRSHHHHHH |

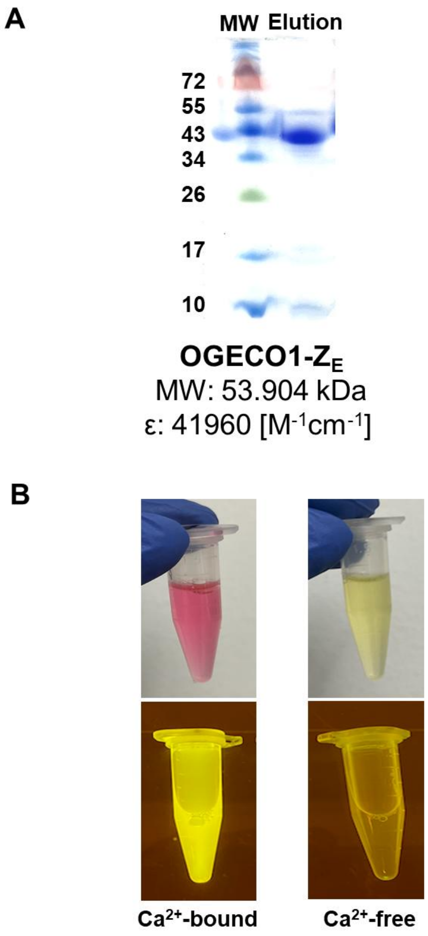


**Figure S1**. (A) The SDS-PAGE gel of purified OGECO1-$Z_E$. The theoretical molecular weights (MW) and extinction coefficients (ε) are provided on bottom of gel image. (B) Photographs of soluble OGECO1-$Z_E$ at the $Ca^{2+}$-free (left) and $Ca^{2+}$-bound state (right), showing changes in color (top) and fluorescence (bottom).

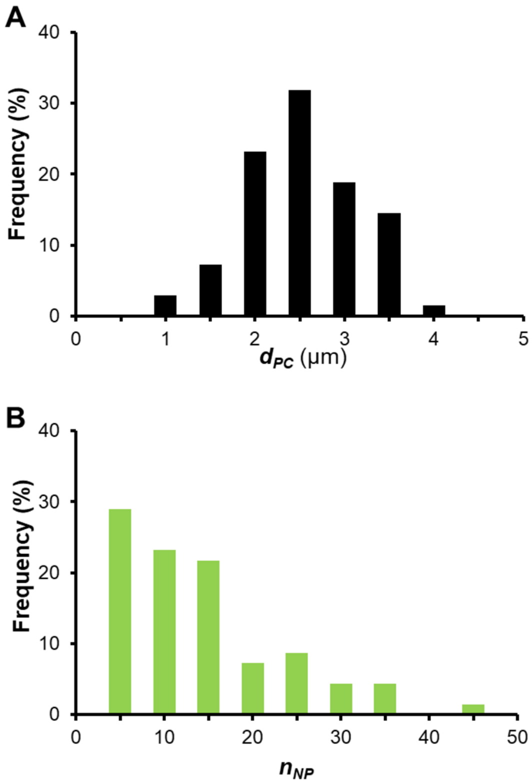


**Figure S2.** (A) Size distribution of mCherry-$Z_E$/$Z_R$-ELP protein cages (PCs) with varying diameters ($d_{PC}$). (B) Number distribution of PCs containing different numbers of encapsulated nanoparticles ($n_{NP}$).

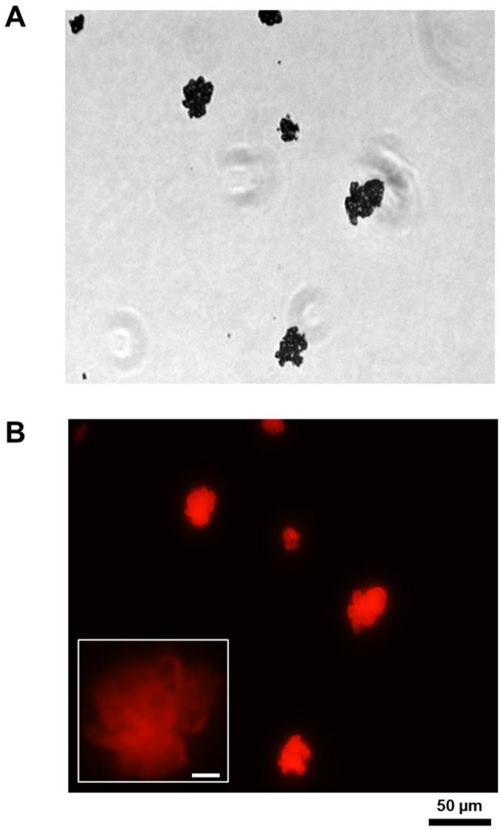


**Figure S3.** Microscope images of mCherry-$Z_E$/$Z_R$-ELP PCs containing $TiO_2$ NPs. (A) Bright-field image. (B) Fluorescence image (Texas Red filter). Inset scale bar: 5µm.

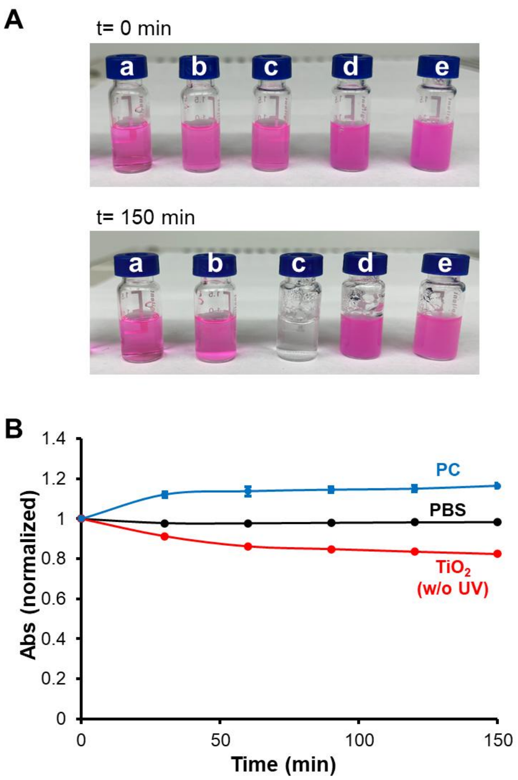


**Figure S4.** (A) Photographs of rhodamine B solutions in phosphate buffered saline (PBS, 2X) taken at 0 min (top) after 150 min exposure to UV light (302 nm). Samples include (a) PBS only, (b) $TiO_2$ NPs without UV exposure, (c) $TiO_2$ NPs, (d) PCs only, and (e) PCs encapsulating $TiO_2$ NPs. (B) Normalized absorbances at wavelength 554 nm as a function of time for samples (a), (b), and (d). Corresponding data for samples (c) and (e) are presented in Figure 5.

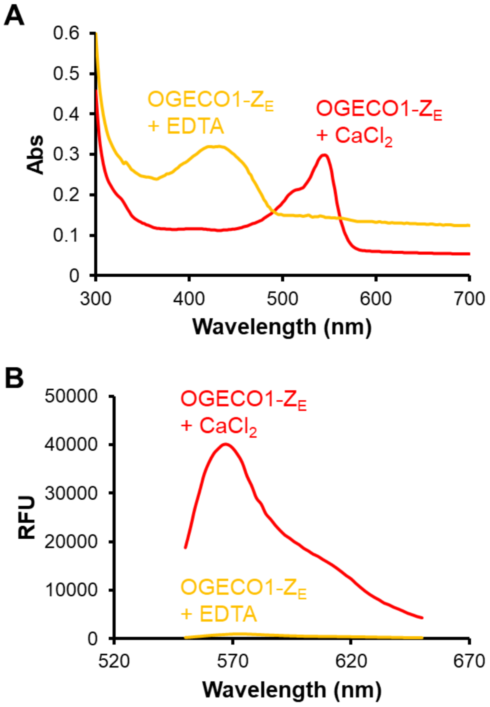


**Figure S5.** (A) Absorbance (Abs) and fluorescence changes of soluble OGEO1-$Z_E$ upon calcium ion binding. (B) Fluorescence emission spectra (relative fluorescence units, RFU) measured with excitation at 520 nm. The spectra were measured in the presence of ethylenediaminetetraacetic acid (EDTA) (yellow) or $CaCl_2$ (red).
